# Regulatory DNA in *A*. *thaliana* can tolerate high levels of sequence divergence

**DOI:** 10.1101/104323

**Authors:** C.M. Alexandre, J.R. Urton, K. Jean-Baptiste, M.W. Dorrity, J.C. Cuperus, A.M. Sullivan, F. Bemm, D. Jolic, A.A. Arsovski, A. Thompson, J.L. Nemhauser, S. Fields, D. Weigel, K.L. Bubb, C. Queitsch

## Abstract

Variation in regulatory DNA is thought to drive evolution. Cross-species comparisons of regulatory DNA have provided evidence for both weak purifying selection and substantial turnover in regulatory regions. However, disruption of transcription factor binding sites can affect the expression of neighboring genes. Thus, the base-pair level functional annotation of regulatory DNA has proven challenging. Here, we explore regulatory DNA variation and its functional consequences in genetically diverse strains of the plant *Arabidopsis thaliana*, which largely maintain the positional homology of regulatory DNA. Using chromatin accessibility to delineate regulatory DNA genome-wide, we find that 15% of approximately 50,000 regulatory sites varied in accessibility among strains. Some of these accessibility differences are associated with extensive underlying sequence variation, encompassing many deletions and dramatically hypervariable sequence. For the majority of such regulatory sites, nearby gene expression was similar, despite this large genetic variation. However, among all regulatory sites, those with both high levels of sequence variation and differential chromatin accessibility are the most likely to reside near genes with differential expression among strains. Unexpectedly, the vast majority of regulatory sites that differed in chromatin accessibility among strains show little variation in the underlying DNA sequence, implicating variation in upstream regulators.

## INTRODUCTION

Changes in gene regulation are thought to be important drivers of phenotypic variation, evolution, and disease. This was first postulated over 50 years ago, when it was reasoned that phenotypic variation among organisms “are the result of changes in the patterns of timing and rate of activity of structural genes rather than of changes in functional properties of the polypeptides as a result of changes in amino acid sequence” (Zuckerkandl and Pauling 1965). Similar conclusions were drawn based on diverse evidence, including the presence of different RNAs in different cell types (Britten and Davidson 1969), the discrepancy between genome size and gene number (Ohno 1972), the discrepancy between high mutation rate and phenotypic robustness (Crow and Kimura 1970; Muller 1966), and the striking similarity of human and chimpanzee proteins (King and Wilson 1975).

‘Active’ genomic regions have been historically identified through sensitivity to endonuclease cleavage (Gottesfeld, Murphy, and Bonner 1975; Weintraub and Groudine 1976; Wu, Wong, and Elgin 1979; Keene et al. 1981; Feng and Villeponteau 1992). DNase I Hypersensitive Sites (DHSs) were often found up- and down-stream of actively transcribed gene bodies (Elgin 1981), demarcating *cis*-regulatory DNA bound by transcription factors (TFs) rather than directly reflecting transcriptional activity (Gross and Garrard 1988). With the advent of high-throughput sequencing, endonuclease hypersensitivity and other genome-scale methods have been used to delineate regulatory DNA in hundreds of human cell types and tissues, plants, fungi, and animals (Sabo et al. 2004; Hesselberth et al. 2009; Furey 2012; Buenrostro et al. 2013; Sullivan et al. 2014; Villar et al. 2015; Weber et al. 2016).

These methods have allowed comparisons of regulatory loci across species on a genome-wide scale. As predicted five decades ago, regulatory DNA generally appears to be under very weak purifying selection – similar to that of four-fold synonymous sites (Stern and Frankel 2013; Vierstra et al. 2014). However, mutation of even a single base pair within a transcription factor binding site can suffice to disrupt regulation of its target gene (Hattori et al. 2002; Liu et al. 2014; Vierstra et al. 2015). Thus, unlike coding regions where the signature of purifying selection predicts gene structure with considerable success, simple conservation metrics largely fail to predict the base-pair-level anatomy of regulatory regions.

Cross-species analyses have revealed that positional conservation of regulatory DNA decays with evolutionary distance, and that, within positionally conserved regulatory sequences, TF binding sites turn over rapidly (J. Kim, He, and Sinha 2009; Shibata et al. 2012; Vierstra et al. 2014; Villar et al. 2015). To make progress towards a base-pair-level functional understanding of regulatory regions, we compared the regulatory regions among less diverged genomes, in which most of their positional homology is maintained and less nucleotide divergence has occurred. The *Arabidopsis thaliana* species offers an excellent set of such genomes: it consists of hundreds of strains that are predominantly self-fertilizing and largely homozygous (Bomblies et al. 2010), with a within-species pairwise divergence similar to the between-species divergence of human and chimpanzee (1001 Genomes Consortium. Electronic address: and 1001 Genomes Consortium 2016), but with enough outcrossing among strains to avoid the effects of Muller’s ratchet (Felsenstein 1974). Linkage disequilibrium decays at a rate similar to that of African humans (S. Kim et al. 2007). Thus, by profiling only a small sample of *A. thaliana* strains, we can capture much of the common within-species variation in regulatory loci and, at the same time, sample many replicate alleles.

Here, we use DNase I-seq (and some ATAC-seq) to map hypersensitive sites (DHSs) for each of five geographically and genetically diverse strains of *A. thaliana*. We compared (i) chromatin accessibility (DHSs) as an indicator of TF-binding; (ii) sequence variation; and (iii) expression levels of nearby genes. Among these strains, we find that 15% of the ~ 50,000 DHSs were differentially accessible. Surprisingly, only a small minority of these differential DHSs were due to underlying sequence variation, putting into question how often variation in regulatory DNA drives phenotypic variation. At the same time, we find instances of dramatic sequence variation in regulatory regions with little or no effect on chromatin accessibility and gene expression. Although chromatin accessibility change *per se* is a poor predictor of gene expression changes in nearby genes (Sullivan et al. 2015), combining changes in chromatin accessibility and underlying DNA sequence variation improved predictions. Taken together, our results illustrate the challenges inherent in understanding regulatory variation and its relationship to phenotypic divergence even in a well-studied model species.

## RESULTS

### Identification of regulatory DNA across five divergent strains

For each of five divergent *A. thaliana* strains (Fig. 1A), we created transgenic lines in which nuclei are biotin-tagged to enable capture with streptavidin beads (Deal and Henikoff 2010; Sullivan et al. 2014). Nuclei from 7-day-old light-grown seedlings for each strain were harvested and treated as described (see Methods) to generate DNase-seq data. DNase I reads of all strains were aligned to the Col-0 reference genome because the quality of genome assemblies were variable. DNase I hypersensitive sites (DHSs) were called as described (Sullivan et al. 2014) (Fig. 1). We and others have found chromatin accessibility profiles to be highly reproducible (Sullivan et al. 2014; Sullivan et al. 2015). Nevertheless, we obtained replicates for two accessions (Col-0, Bur-0), using an alternative method to assess chromatin accessibility, ATAC-seq. As expected, DHSs were highly reproducible across replicates and methods (Supplemental Fig. S1).

**Figure 1.**
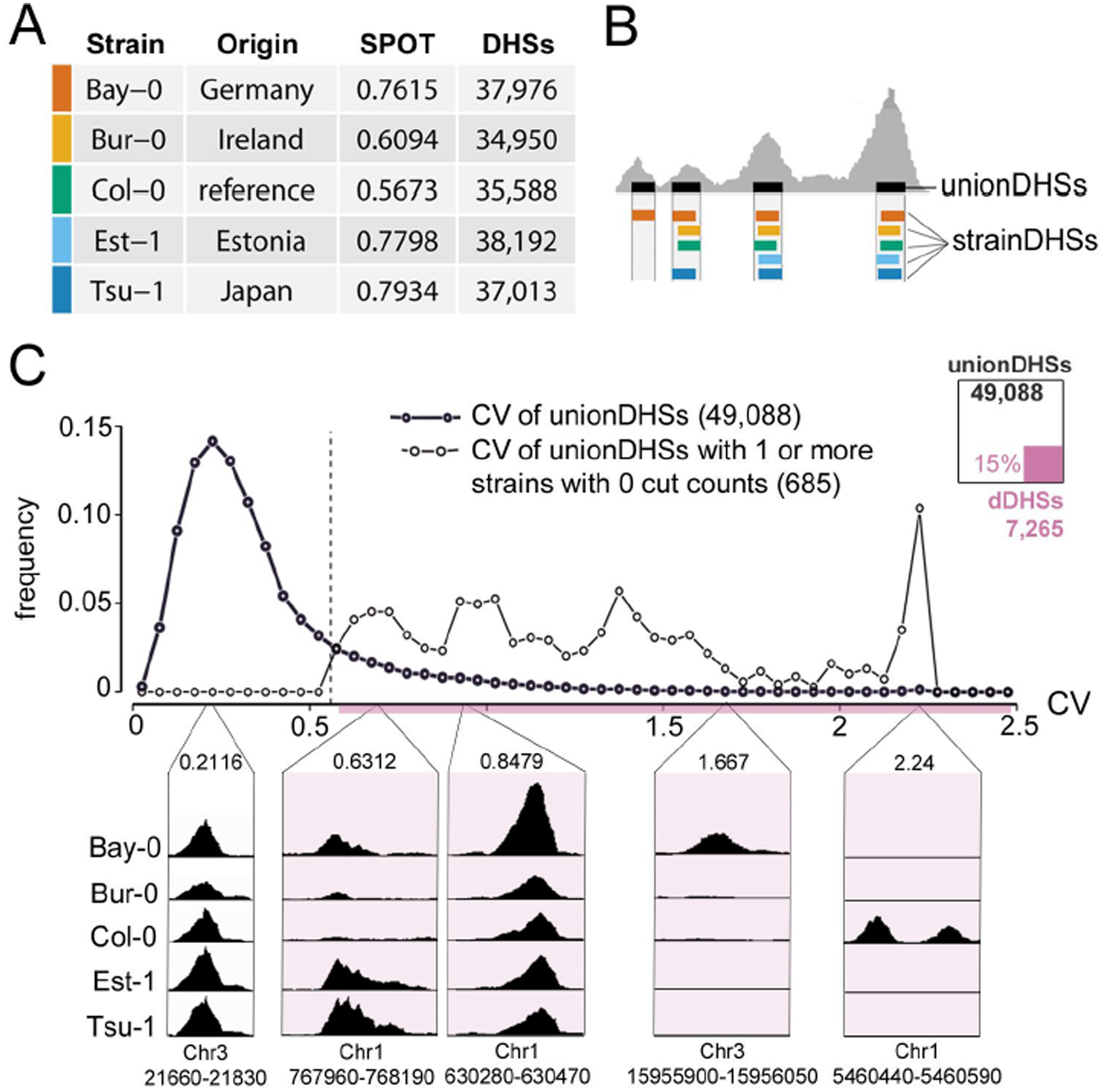
Identifying regions of differential chromatin accessibility among five *A. thaliana* strains. (*A*) Data and data quality for the five strains examined. SPOT score, a metric of data quality, describes fraction of cuts within hotspots (Sullivan et al. 2015). Colors indicate specific strains throughout manuscript (palette is colorblind-accessible (Wong 2011); Bay-0, vermillion; Bur-0, orange; Col-0, bluish green; Est-1, sky blue; Tsu-1, blue; differential DHSs, reddish purple). (*B*) Schematic depiction of deriving union DHSs (uDHSs) for subsequent analysis. (*C*) Distribution of coefficient of variation (CV) among union DHSs (bold line). Distribution of coefficient of variation (CV) among the subset of union DHSs for which at least one strain has no cut counts (lighter line). A CV threshold of 0.56 was chosen to include all uDHSs in the latter category. This approach categorized 15% of all union DHSs as differentially accessible (see inset diagram, dDHSs in purple). See Supplemental Fig. S2 for further details on the rationale for CV as a metric for identifying dDHSs. Examples of individual uDHSs with respective chromosome coordinates and CVs are shown below CV distribution.

We merged DHSs from the five strains to generate a union set of 49,088 DHSs (Fig. 1B, Supplemental Table S1). Most union DHSs were shared among the five strains, providing yet another means of replication; only 8,778 DHSs were private, *i.e.* called in only one strain. Private DHSs tended to show comparatively low nuclease cut counts, on the edge of our detection criteria, and therefore were potentially enriched for false positive DHSs (Supplemental Fig. S2A, B). Therefore, rather than calling DHS presence or absence across strains, we compared DNase I cut counts within union DHSs to identify differential DHSs among strains.

Specifically, because DNase I cut count standard deviation (σ) and mean (μ) showed an approximately linear relationship across the five strains (Supplemental Fig. S2C), we used a coefficient of variation (CV, σ/μ) threshold to define 7,265 DHSs (15%) as differentially accessible DHSs (differential DHSs, dDHSs, Fig. 1C, see Methods).

### Reference bias arises from substantial structural variation in regulatory loci

A majority (62%) of the 7,265 differential DHSs (n=4,508) were most accessible in the reference strain Col-0 (dDHSs-C) (Fig. 2A), presumably because the reference Col-0 genome was used for alignments. The remaining differential DHSs (n=2,757) were most accessible in each of the other four strains at approximately equal numbers (see Fig. 2A, inset). Differential DHSs tended to show a bimodal distribution of accessibility (Supplemental Fig. S2D), and for the differential DHSs in which only one strain displayed considerably different cut count (74%, 3,775 of 5,131 dDHSs), the outlier strain was Col-0 with highest accessibility.

**Figure 2.**
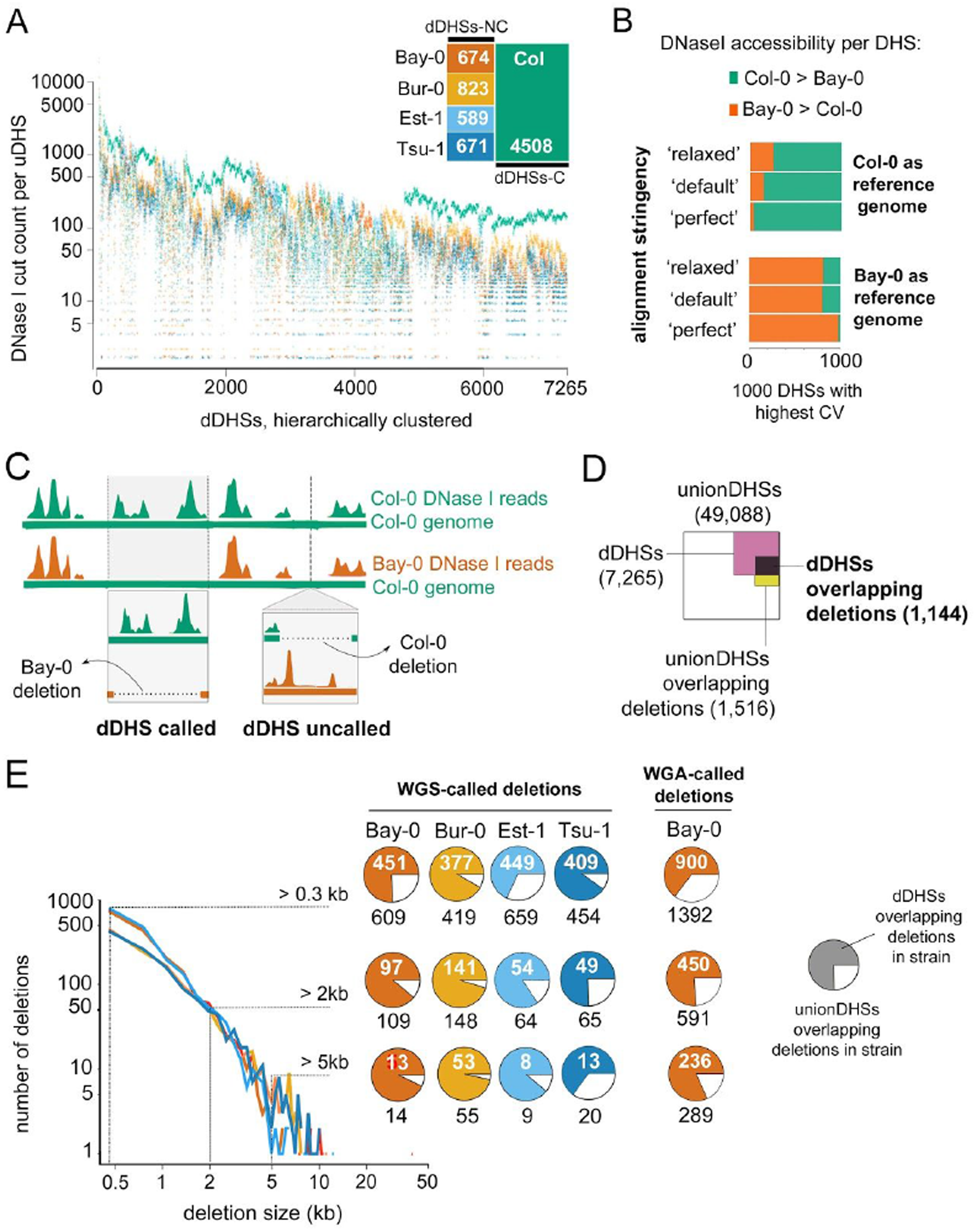
Structural variation contributes to profound reference bias and explains a sizable minority of differential DHSs. (*A*) DNase I-cut counts for each of 7,265 dDHS across all five strains (color-coded dots, see inset) are shown. dDHSs are displayed hierarchically clustered with each column representing a given DHS. dDHSs in which the reference Col-0 (green) shows more cut counts than any of the other strains (dDHSs-C) are most common. (*B*) The reference bias in (*A*) arises through use of the Col-0 reference genome for DNase I read alignment. Two different reference genomes, Col-0 (green, top) and Bay-0 (vermillion, bottom), were compared for alignment at different stringencies. Replacing the Col-0 reference genome with the Bay-0 draft genome for read alignments, resulted in a Bay-0 specific bias, *i.e.* now the majority of dDHSs was most accessible in Bay-0. The 1000 dDHSs with the highest CV were used for analysis. Requiring perfect alignment (no mismatches) between DNase I read and reference genome enhanced this effect, while relaxing alignment parameters dampened this effect in the Sanger-sequenced Col-0 genome. (see Methods for details). (*C*) Schematic outlining the consequences of missing sequence in Col-0, the reference strain, versus missing sequence in a non-Col-0 strain for bias in calling dDHSs. In our analysis, DNase I reads from each strain were aligned to the Col-0 reference sequence (depicted by green horizontal bar). If a Col-0 sequence corresponding to a DHS was missing in Bay-0, this DHS would appear inaccessible in Bay-0 and called as a dDHS-C. However if a Bay-0 sequence corresponding to a DHS was missing in Col-0, the DHS would not be included in the union DHS set and not be counted as a dDHS-NC. (*D*) Overlap between WGS-called deletions, union DHSs and dDHSs shows that differential DHSs are enriched for strain-specific deletions; areas are proportional to the size of each category. (*E*) Size distribution of WGS (whole-genome shotgun sequence)-called deletions in each strain was similar, with few deletions over 20 kb. Pie charts indicate the fraction of union DHSs overlapping strain-specific WGS-called deletions of various minimum sizes that were characterized as differential DHSs (see adjacent key). For comparison, see the fraction of union DHSs overlapping Bay-0 deletions of various minimum sizes called by WGA (whole-genome alignment) of the Bay-0 draft genome to the Col-0 reference genome (see Methods).

Approximately 4% of any *A. thaliana* strain genome is estimated to be absent in another strain (Clark et al. 2007; Plantegenet et al. 2009). Compared to coding DNA, regulatory DNA may be disproportionately affected by strain-specific variation because it is accessible and less constrained by selection. Differential DHSs can arise in three ways: through (i) strain-specific deletions of the sequence underlying a DHS; (ii) strain-specific variation in the sequence underlying a DHS that affects TF binding (both *cis*-effects); and (iii) by strain-specific variation in activity or DNA binding preference of transcription factors (*trans*-effects). To explore the relative contributions of these three categories to the differential DHSs, we had to resolve the cause and extent of the reference bias, likely due to the use of the Col-0 reference genome for alignments.

To test this assumption, we obtained a *de novo* Bay-0 genome assembly. Aligning both Bay-0 and Col-0 DNase I reads to this Bay-0 reference reversed the previously observed bias, such that the majority of differential DHSs were now most accessible in Bay-0 (Fig. 2B, C). To explore sequence divergence at the base-pair level as a cause of reference bias, we used three different alignment stringencies, aligning to both the Col-0 and Bay-0 reference genomes. Consistent with the effects of sequence divergence, the reference bias was exaggerated when we required perfect alignment between sequence read and reference genome; conversely, the reference bias was diminished when the alignment requirements were relaxed (Fig. 2B). The reference bias was not diminished with relaxed alignment to the Bay-0 genome, presumably because it is not of the same quality as the Col-0 reference genome (Fig. 2B). In summary, our analysis demonstrates that there is no true excess of differential DHSs that are most accessible in Col-0; rather, due to using the Col-0 genome for alignment, we undercounted differential DHSs that are accessible in the other strains.

Having established the validity of differential DHSs, we explored whether they differed in genomic or biological features from union DHSs from which they are drawn. Compared to union DHSs, differential DHSs were more likely to reside in coding regions and transposons and less likely to reside in intergenic regions (Supplemental Fig. S3A). As sequence variation is implicated in generating accessibility difference, we compared recombination rates within union DHSs, differential DHSs, and comparable regions outside of DHSs. Using recombination events imputed from the mosaic genomes of the MAGIC lines (Kover et al. 2009), a higher fraction of DHSs (2.1%) overlapped with a recombination event compared to same-length regions shifted 5 and 10 kb (1.7%, p<10-9), consistent with recombination favoring accessible regions in plants (Rodgers-Melnick et al. 2016). Focusing on recurrent recombination events, 9% of differential DHSs overlapped with recombination events in multiple lines compared to only 3.8% of union DHSs. Comparing gene annotations for genes residing either near union or differential DHSs, we found that the latter were enriched for annotations related to programmed cell death (Supplemental Fig. S3B). The features of differential DHSs (location, elevated recombination rate, and proximity to defense-and cell death-related genes) are consistent with prior work on defense-related genes, recombination, and regulatory sequence variation (Choi et al. 2016; Gan et al. 2011), and together with the observed reference bias, suggest that a substantial fraction of differential DHSs may be due to the absence of DHS sequence (a local deletion) in one or more strain.

In the absence of comparable genome assemblies for the non-Col-0 strains, we used available short read data (1001 Genomes Consortium. Electronic address: and 1001 Genomes Consortium 2016) to call putative deletions in Bay-0, Bur-0, Est-1 and Tsu-1 (Supplemental Tables S2-5) to determine the fraction of union DHSs that only appear differentially accessible due to sequence divergence, rather than to being in fact differentially accessible. For comparison, we also conducted this analysis for the Bay-0 short-read data and the novel Bay-0 assembly (Fig. 2E). Specifically, we used whole-genome shotgun (WGS) read coverage to predict putative deletions of 300 bp or greater in each strain, including Bay-0, with respect to the Col-0 genome (Fig. 2E, see Methods). The size distribution of predicted deletions among strains was similar, with some deletions extending over 40 kb. In total, of the 49,088 union DHSs, 1,516 DHSs (3%) overlapped with a predicted deletion by at least one bp; of the 7,265 differential DHSs, 1,144 DHSs (16%) overlapped with a predicted deletion (Fig. 2D). We reasoned that strain-specific deletions overlapping with union DHSs should result in strain-specific differential DHSs. Indeed, in particular with increasing deletion size, we observed that increasing numbers of union DHSs overlapping with predicted deletions were called as differential DHSs (Fig. 2E). However, the overlap of predicted deletions with differential DHSs was imperfect, either due to miscalling predicted deletions or miscalling differential DHSs. We assessed the accuracy of our deletion predictions by comparing our results to two independent data sets: the Bay-0 comparative genomic hybridization (CGH) array data (Plantegenet et al. 2009) and the novel Bay-0 genome assembly. We found strong correspondence of our predicted deletions with both data sets (Supplemental Fig. S4). Moreover, deletions called by whole genome alignment (WGA-called deletions; see Methods) behaved like deletions called by read coverage for Bay-0 WGS data (Fig. 2E).

### Sanger-sequencing of wrongly predicted deletions reveals hypervariable DHS sequences

To directly assess the accuracy of our predictions, we PCR-amplified regions corresponding to ten WGS-called deleted DHSs that reside near well-annotated genes. Of these ten regions, seven were indeed deleted (Fig. 3A, Supplemental Table S6). For the remaining three, Sanger-sequencing revealed that strains predicted to have deletions carry instead a homozygous DHS allele with dramatically different sequence but of approximately equal length as the Col-0 DHS allele (Fig. 3B). We Sanger-sequenced eight additional strains and found that each strain was homozygous for one of the two previously-identified alleles, consistent with a single mutation event creating the vast majority of the nucleotide differences in each of these regions (Fig. 3B, Supplemental Fig. S5).

**Figure 3.**
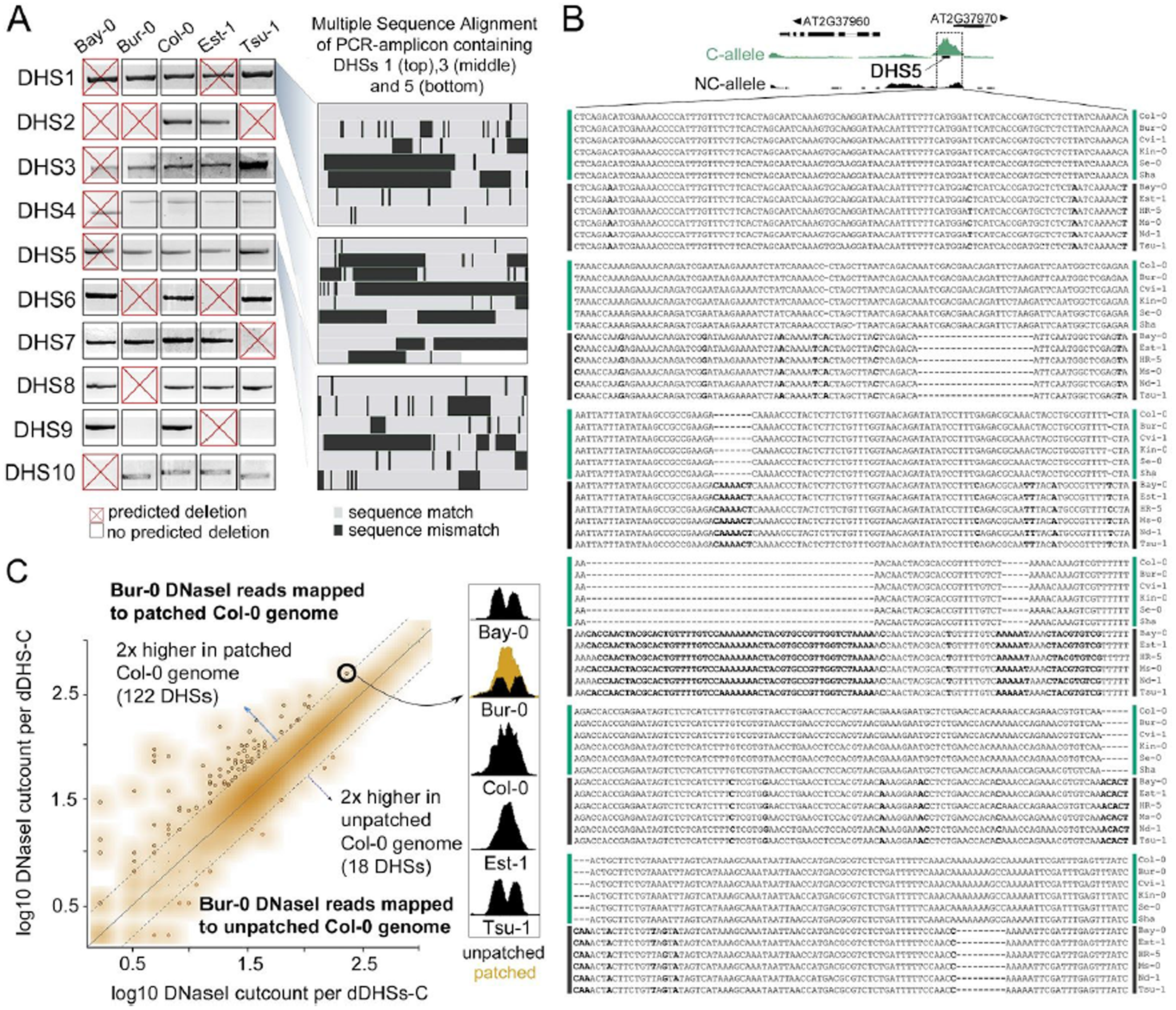
Hypervariable sequence coinciding with DHSs poses a challenge for sequence alignment but does not significantly contribute to reference bias. (*A*) PCR confirmed seven out of ten predicted deletions in at least one strain through decreased size or absence of diagnostic PCR product. Predicted deletions are denoted as red X. In some instances, strains carried a deletion that was not predicted (black box, DHS9 in Bur-0 and Tsu-1, false negatives). For the three wrongly predicted deletions coinciding with DHS1, DHS3, and DHS5, Sanger-sequenced PCR-products from 12 *A. thaliana* strains revealed that strains carried either a homozygous Col-like sequence allele or a homozygous non-Col-like allele with dramatically different sequence but approximately equal length. Alignments are represented as thumbnails with 100 bp per line, identical sequence in grey and mismatches in black. Col-0 coordinates for these three hypervariable loci are Chr1:5,673,357-5,674,171; Chr1:28,422,395-28,423,622; and Chr2:15,890,369-15,891,184, respectively. (*B*) For DHS5, the base-pair resolution multiple sequence alignment across 12 strains is shown with mismatches highlighted and allele-type indicated (Col-like allele, C-allele, green; non-Col-like allele, NC-allele, black). Above, DHS5 screenshot with dramatically lower cut counts in strains with NC-allele (black). For base-pair resolution multiple alignments of DHS1 and DHS3, see Supplemental Fig. S5A, B. (*C*), Scatterplot of Bur-0 DNase I reads aligned to either patched or unpatched Col-0 sequence. The patched Col-0 genome was generated by replacing DHS-sequences with locally-assembled sequence from Bur-0 WGS reads (see Methods). Dotted lines indicate 2-fold higher cut counts for respective DHSs with either patched or unpatched genome. The majority of DHSs was not affected (orange cloud). At right, an example of false positive differential DHS for which patching resulted in higher numbers of aligned DNase I reads such that ‘patched’ cut count approximated that of the Col-0 DHS. The similarity of DHS pattern between Bur-0, Bay-0 and Tsu-1 suggests that the differential DHS in the two latter strains is also a false positive due to sequence hypervariability.

We refer to DHSs with two highly disparate alleles as hypervariable DHSs (hypDHSs). The existence of hypervariable DHSs suggested that, in addition to undercounting differential DHSs most accessible in non-Col strains (due to the absence of DHS sequences in Col-0), we may be overcounting differential DHS most accessible in Col-0. Neither read-coverage-based detection of deletions nor comparative genomic hybridization data can distinguish between true deletions and sequence regions that are diverged to the point of precluding alignment of short reads.

Therefore, we explored the extent of DHS hypervariability by Sanger-sequencing ten additional DHSs with very high CV across the five strains. We found elevated sequence diversity in all ten sequence regions compared to the genome-wide background; the mean number of substitutions, deletions and insertions (SDI) was 14.3 across the 150 bp DHS regions. The immediate area surrounding these DHSs (~800 bp total, including DHS) also showed elevated sequence diversity (Supplemental Table S7).

Next, we aimed to recover the true sequence of all differential DHSs that were most accessible in Col-0 to determine to which extent hypervariability affected our analysis. We used PHRAP (Phil Green, personal communication) to perform local *de novo* assemblies of the Bur-0 DHS allele for each of the 4,508 differential DHS most accessible in Col-0, using (i) a Col-0 backbone sequence with per-bp quality set to zero, and (ii) all pairs of WGS reads in which at least one read mapped to that DHS region, *i.e.* using one of the paired reads to anchor the other, even if the sequence difference from the Col-0 allele prohibited alignment (see Methods). We tested this strategy by comparing our local assemblies to the Bur-0 Sanger sequence for the ten DHSs, in addition to available Bur-0 genome sequence (Gan et al. 2011). Comparing patched sequence to respective Sanger-sequence, across the ten DHSs, 83% of SDIs were corrected (120/143); for six DHS, sequences were completely corrected. Eleven of the 23 uncorrected SDIs were indels. Comparing the published Bur-0 assembly to the respective Bur-0 Sanger sequences, only seven of the143 SDIs were missed, all of which were substitutions clustered within a single DHS (Supplemental Table S7).

As patching proved effective, we patched the Col-0 genome with the *de novo* derived Bur-0-like sequences, maintaining genome coordinates, and then re-aligned the Bur-0 DNase I reads. For 122 of the 4,508 differential DHSs-C, we aligned more than twice as many Bur-0 DNase I reads to the patched Col-0 genome compared to the reference Col-0 genome (Fig. 3C). However, perhaps surprisingly, the majority of previously-called differential DHSs-C loci remained most accessible in Col-0. Moreover, when we recalculated CV across all strains using the newly patched genome for Bur-0, only 100 DHSs out of 4,508 differential DHSs-C fell below the differential DHSs threshold (Fig. 1C), with most (84/100) close to the threshold (Fig. 1C). In summary, the observed excess of differential DHSs most accessible in Col-0 was largely due to undercounting in other strains (false negatives), and not to poor alignment of strain-specific DNase I reads to the Col-0 reference genome (false positives).

### Hypervariable differential DHSs tend to reside near differentially expressed genes

The DHS landscape for a given genotype is largely static, with only 5-10% of DHSs changing in accessibility in pairwise comparisons of conditions (Sullivan et al. 2014) or development stages (Sullivan et al., *in review*) in spite of widespread changes in gene expression. Static DHS presence is generally not a good predictor of nearby gene expression for multiple reasons; for example, accessibility can be caused by repressors and DHS assignment to nearby genes is an oversimplification. However, the presence of dynamic DHSs more frequently correlates with expression changes in nearby genes (Sullivan et al. 2014; Sullivan et al. 2015).

Across genotypes, we had to consider the additional complexity of strain-specific sequence variation underlying DHSs. Thus, we explored the relationship of high sequence variation with both DNase I accessibility and expression of nearby genes. As previously observed (Plantegenet et al. 2009), deleted regulatory regions often reside near deleted genes, providing a trivial reason for differential gene expression unrelated to the evolution of regulatory DNA. Therefore, to select for deleted DHSs near non-deleted genes, we used a set of smaller predicted deletions, generated by merging only overlapping 150 bp windows of low WGS coverage and retaining all predicted deletion windows, not only those >300 bp as before (Supplemental Table S8). Of these smaller predicted deletions (>=150 bp), 308 overlapped with a union DHS but not a gene. Comparable, high-quality expression data were publicly available for Bur-0 and Col-0 strains (Gan et al. 2011); hence, we considered these strains for further analysis. We applied DESeq (Anders and Huber 2010) to these data (Gan et al. 2011) and identified 737 protein-coding genes with differential expression between Bur-0 and Col-0.

We then determined for Col-0 and Bur-0 the proximity of the various DHS sets (Bur-0/Col-0 union DHSs, uDHSs; differential DHSs, dDHSs; deleted DHSs, delDHSs, hypervariable DHSs, hypDHSs) to the nearest differentially expressed gene, assessing DHSs residing upstream and downstream of transcription start sites. Only a small fraction of Bur-0/Col-0 union DHSs (Supplemental Table S9) resided near differentially expressed genes (Fig. 4A; for specific examples see Fig. 5A, top). As expected, this fraction increased, albeit only modestly, when differentially accessible DHSs were considered. HypDHSs were more likely than dDHSs to reside near differentially expressed genes, regardless of whether they differed in accessibility. However, hypDHSs with altered accessibility predicted expression changes in nearby genes almost as well as delDHSs. Taking into account DHSs residing downstream of transcription start sites, delDHSs predicted expression changes for approximately 25% of nearby genes.

**Figure 4.**
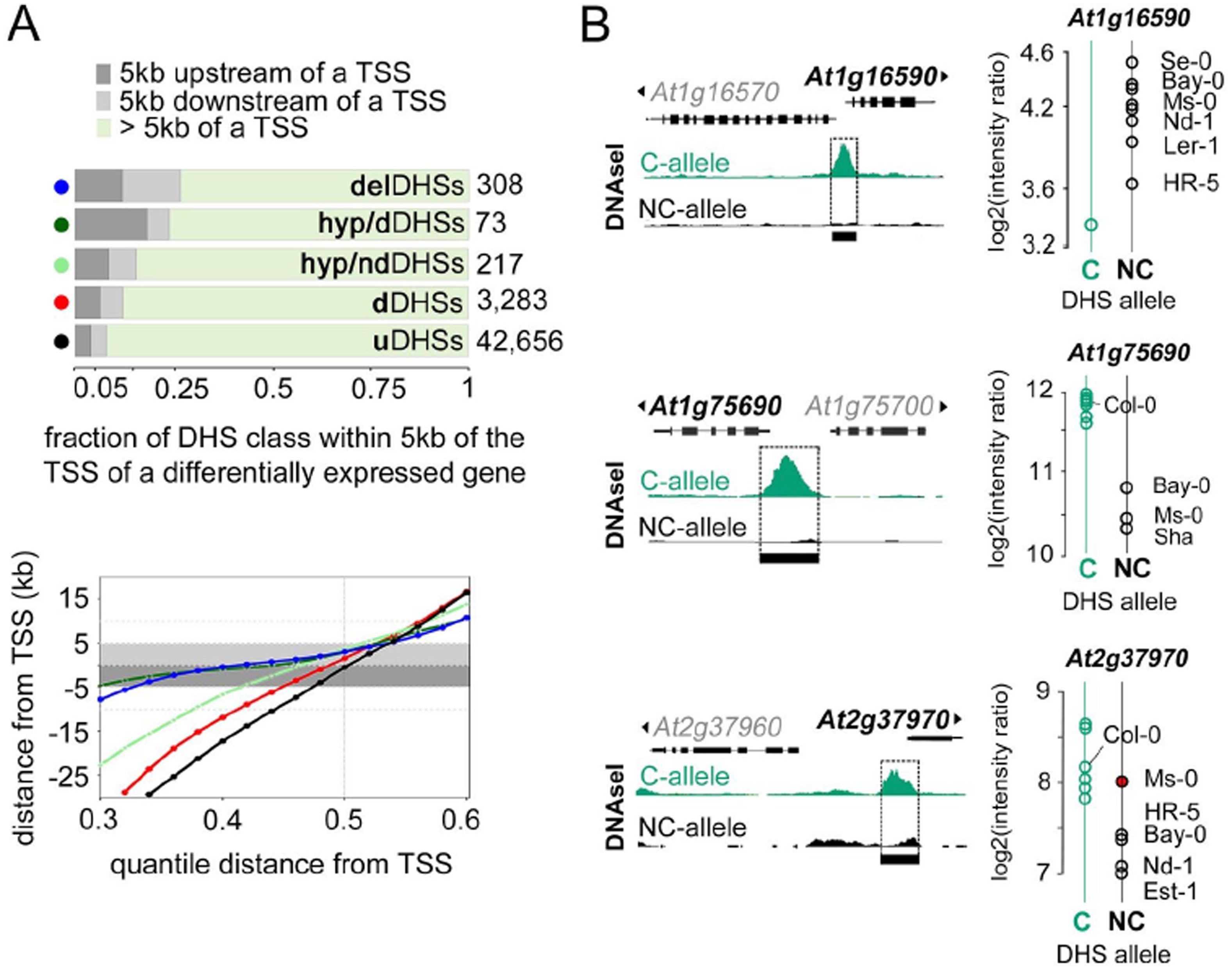
DHSs alleles with high sequence divergence tend to reside near genes with different expression levels. (*A, Top*) Fraction of DHSs residing within 5 kb of a differentially expressed gene by DHS sets (color-coded). DHSs were subsampled to sets of 50 to allow comparisons among the examined DHSs sets, which occurred at vastly different frequency (see Methods). Deleted DHSs (blue circle) and hypervariable DHSs (hypDHSs, light and dark green circles) — particularly those in which accessibility also differed (hyp/d DHSs, dark green circles) — were more likely to reside near differentially-expressed genes than either dDHSs (red circles) or union DHSs (black circles). Differential accessibility was observed for 73 of 290 of hypervariable DHSs (hyp/d DHSs, dark green circles). (*A, Bottom*) Distributions of distances between DHSs sets (color-coded as above) to TSS of a nearby differentially-expressed gene (see Methods). (*B*) For the three previously-identified hypervariable DHSs (DHS1, DHS3 and DHS5, Fig. 3A, B), the DHS-allele (Col-like C-allele in green, Non-Col-like NC-allele in black) determined by Sanger-sequencing predicted the expression level of a neighboring gene in eleven strains with publicly-available expression data (Lempe et al. 2005) with one exception (Ms-0 in DHS5, red dot). At left, screenshots showing altered DHS accessibility at these sites (see also Fig. 3B); at right, gene expression (as log2(intensity ratio), Lempe et al. 2005) for strains carrying either the DHS C-allele (green) or the DHS NC-allele (black).

In summary, neither sequence variation nor accessibility variation were good predictors of expression changes in nearby genes; however, combining this information allowed far better predictions than relying on either factor alone (Fig. 4A, B). For example, for the three original hypDHSs — originally mistaken for deletions — we found that, for each locus, allele type corresponded to expression level of nearby gene for 11 strains with publicly available expression data (Lempe et al. 2005)
(Fig. 4B).

These three hypDHSs reside near well-annotated genes, offering the opportunity to examine whether differential hypDHS alleles are associated with phenotypic variation in addition to expression variation. We focused on the hypDHS upstream of *REV7/*AT1G16590, a gene associated with UV-tolerance in different organisms (Broomfield, Hryciw, and Xiao 2001). *A. thaliana rev7* mutants show moderately increased sensitivity to prolonged UV exposures (Takahashi et al. 2005). *REV7* is expressed at lower levels in Col-0 than Bur-0 and other strains containing the non-Col-like allele (Fig. 4B, top); however, Col-0 and Bur-0 differed only subtly in their growth response to UV exposure (Supplemental Fig. S6).

### Differential DHSs may be conditionally important

Differential DHSs are thought to be most informative for understanding the regulatory dynamics underlying specific conditional responses and developmental trajectories. For a single expression data set from 11-day-old seedlings grown in long days (Gan et al. 2011), only 25% of delDHSs resided near differentially expressed genes (*e.g.* Fig. 5A, top, Fig. 4A), suggesting that the remaining 75% of delDHSs (*e.g.* Fig. 5A, bottom, Fig. 4A) may matter under different conditions. Accessibility, especially at conditional regulatory sites, often reflects a poised state with bound but not fully activated transcription factors (Gross and Garrard 1988; Zentner, Tesar, and Scacheri 2011; Nelson and Wardle 2013). We noted that genes implicated in response to various treatments (cold, growth in the dark, pathogen response) often resided near small deletions affecting one of multiple adjacent DHSs, consistent with the idea of conditional regulation occurring at different or combined DHS sites. Therefore, we systematically examined whether conditionally expressed genes in Col-0 tend to reside near multiple DHSs (see Methods).

**Figure 5.**
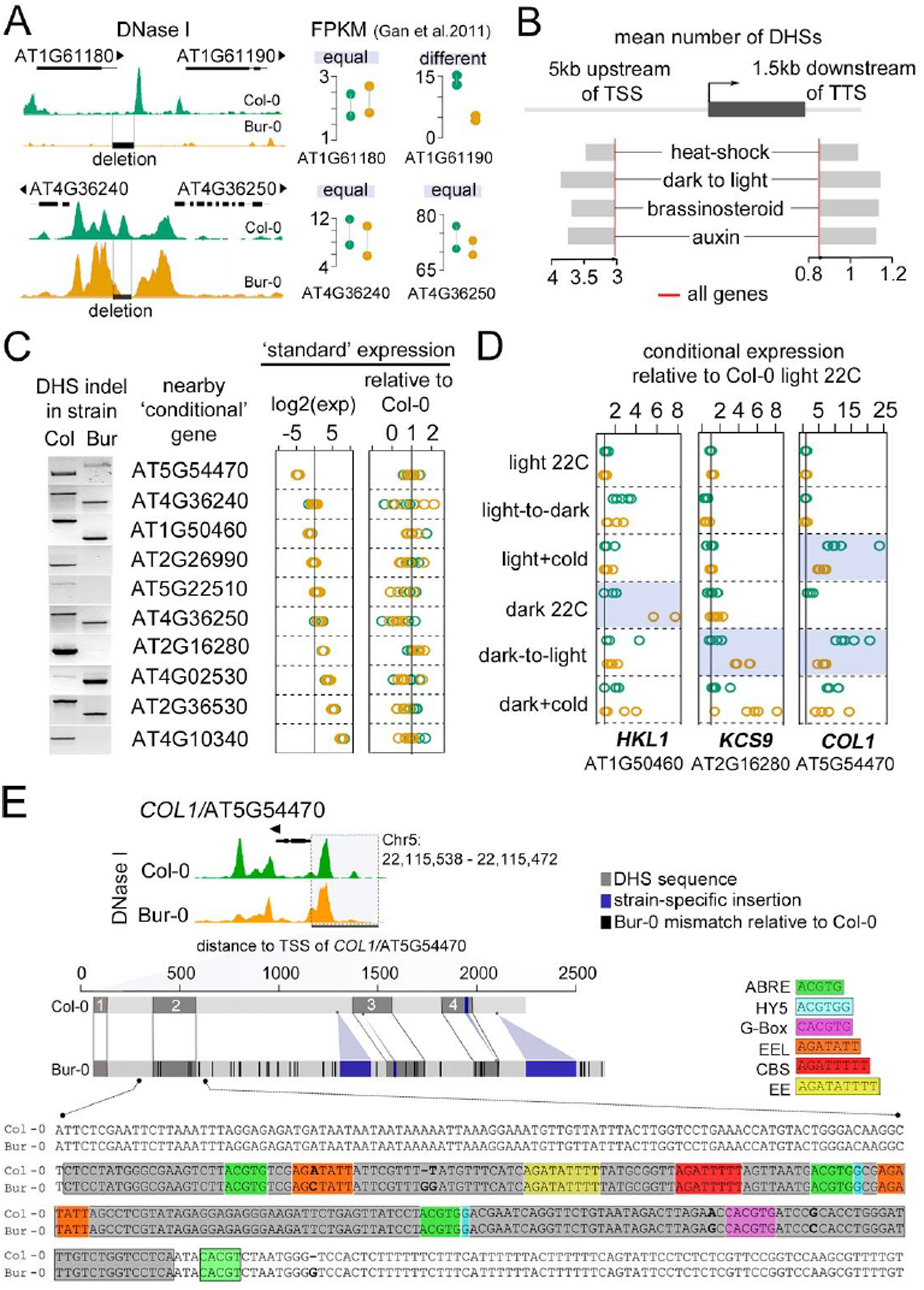
Altered DHS accessibility among strains may have condition-dependent effects on expression of nearby genes. (*A, Top*) A DHS deletion (gray box, borders outlined in black) in Bur-0 resides near a likely target gene with vastly different expression in Bur-0 (orange) compared to Col-0 (green, expression in FPKM) (Gan et al. 2011). (*A, Bottom*) A DHS deletion in Bur-0 (denoted as above) resides near a putative target gene with similar (low) expression levels in Bur-0 and Col-0. (*B*)On average, genes with dynamic expression in response to heat-shock, dark-to-light transition, brassinosteroid and auxin treatment are flanked by significantly more DHSs than all genes (indicated by red line; *i.e.* on average, ~3 DHSs reside 5kb upstream of a gene). DHSs both upstream (left) and downstream (right) showed this tendency. (*C, Left*) Gel images confirming DHS deletions near conditionally-expressed genes. The first example represents an insertion in Bur-0 rather than a deletion. (*C, Middle*) qPCR expression analysis for these genes (*left*) confirms their expression rank order previously observed by RNA-seq (Gan et al. 2011) (*right*) that under standard growth conditions, conditionally-expressed genes were not differentially expressed between Bur-0 (orange) and Col-0 (green). (*D*) Evidence of strain-specific conditional expression (highlighted in blue) associated with a DHS deletion or insertion near *HKL1*/AT1G50460, *KCS9*/AT2G16280, and *COL1*/AT5G54470. (*E, Top*) DNase I screenshot for the region containing *COL1*/AT5G54470 in Col-0 and Bur-0. (*E, Middle*) Schematic alignment of the 2.5 kb region upstream of *COL1*/AT5G54470 with dark gray boxes denoting DHSs, dark blue denoting strain-specific insertions and black lines denoting single nucleotide differences in Bur-0 compared to Col-0. (*E, Bottom*) Base-pair alignment of a 398 bp window (Chr5:22,115,738 - 22,116,135) containing the second and most accessible DHS upstream of *COL1*/AT5G54470. DHS (gray box), transcription factor motifs (color boxes, see key) and indels and/or mismatches between Col-0 and Bur-0 (bold letters) are indicated. ABRE and EEL motifs have been implicated in plant response to cold (Mikkelsen and Thomashow 2009).

Indeed, conditionally expressed genes showed significantly more DHSs both 5 kb upstream of their transcription start site and 1.5 kb downstream of their transcription termination site (Fig. 5B). To test whether some of these DHSs mattered under non-standard growth conditions, we examined DHSs predicted to be deleted, the most extreme form of sequence divergence, for cases in which (i) expression of both neighboring genes was the same in Col-0 and Bur-0 in standard growth conditions (Gan et al. 2011)
, and (ii) expression of a neighboring gene(s) changed in response to various treatments, prioritizing readily testable conditions.

We identified ten genes near such DHSs with prior evidence for conditional expression in cold and/or growth in the dark, but similar expression levels in Bur-0 and Col-0 under standard growth conditions (Fig. 5C, Supplemental Table S10). We PCR-confirmed the predicted DHSs deletions; in nine instances a deletion resided in Bur-0, and in one case DHS sequence was absent in Col-0 (Fig. 5C). We then examined whether any of these ten genes differed in expression between Bur-0 and Col-0 in response to test conditions (see Methods, Fig. 5D, Supplemental Fig. S7). For three of these genes, expression levels differed between Bur-0 and Col-0 in at least one test condition (Fig. 5D). For example, *COL1* expression increased significantly more in Col-0 than in Bur-0 upon dark-to-light transition and in the cold (Fig. 5D, Supplemental Fig. S7). *COL1* expression is known to increase upon exposure to light (Soitamo et al. 2008) and cold (Mikkelsen and Thomashow 2009). Almost all cold-responsive motifs (Mikkelsen and Thomashow, 2009) are contained within the most accessible DHS in the *COL1* regulatory region (Fig. 5E, Supplemental Fig. S8). To explore possible sequence differences between Bur-0 and Col-0 in the extended *COL1* regulatory region, we Sanger-sequenced over 2 kb upstream of *COL1*, which included four union DHSs and the predicted insertion (Fig. 5E, Supplemental Fig. S8). Within the most accessible DHS, we found five single base pair differences between Col-0 and Bur-0, one of which disrupted a cold-response motif. We also discovered an additional 152 bp insertion in Bur-0 between the second and third DHSs which affects spacing of regulatory motifs, as well as smaller insertions, deletions, and multiple single base pair changes (Fig. 5E, Supplemental Fig. S8). Previous large-scale genomics efforts failed to detect these sequence differences. Any one of these regulatory sequence variants or combinations thereof may cause the observed differences in conditional expression. For detailed information on the other two loci with condition-specific effects see Supplemental Text.

### The prevalence of differential DHSs in the absence of sequence variation implies a considerable role of *trans*-effects

We found a surprisingly large proportion of differential DHSs with little strain-specific sequence variation. In the new Bay-0 assembly, 84% and 81% of union and differential DHSs, respectively, carry two or fewer SDIs (Fig. 6A). The mean SDI per DHS was higher in differential DHSs than union DHSs (2.7 compared to 1.8), but this difference was largely driven by outliers (Fig. 6A).

**Figure 6.**
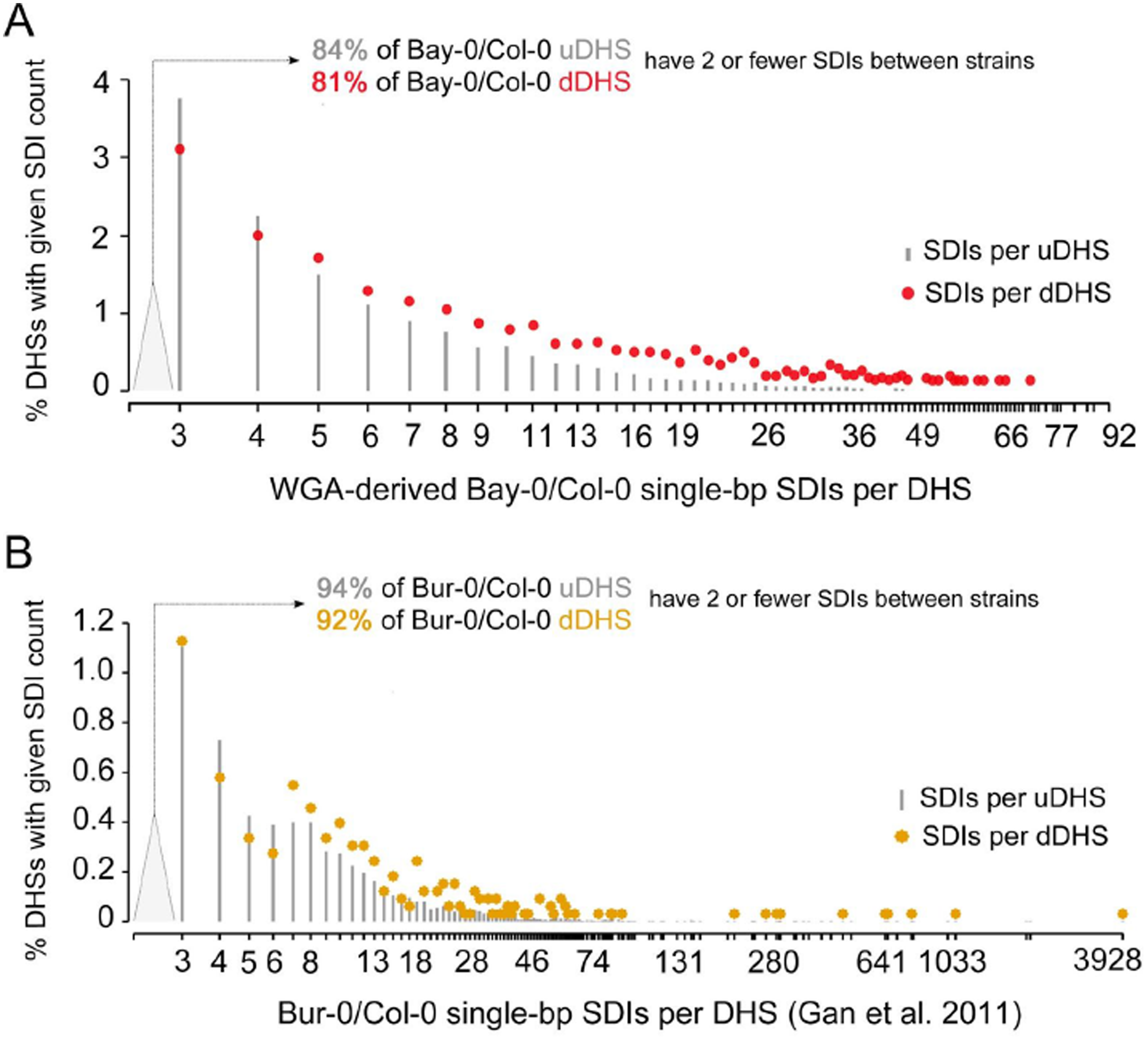
Most differential DHSs are not explained by sequence divergence. (*A*) Total single nucleotide Substitutions, Deletions or Insertions (SDIs) per DHS between Bay-0 and Col-0 were derived from whole genome alignment of the Bay-0 draft genome sequence and Col-0 reference genome sequence (see Methods). X-axis denotes number of SDI per union (gray lines) and dDHS (red dots). Y-axis denotes percentage of DHSs with given number of SDI. (*B*) Total single nucleotide Substitutions, Deletions or Insertions (SDIs) per DHS between Bur-0 and Col-0 (Gan et al. 2011). X-axis denotes number of SDI per union (gray lines) and dDHS (orange dots). Y-axis denotes percentage of DHSs with given number of SDI.

Similarly, in the published Bur-0 sequence, 94% and 92% of union and differential DHSs, respectively, carry two or fewer SDIs (Fig. 6B). As with the Bay-0 genome, the higher mean SDI in Bur-0 dDHSs compared to union DHSs (3.8 versus 1.2) was driven by high-SDI outliers (Fig. 6B). The differences in SDI values for Bay-0 and Bur-0 genomes in comparison to the Col-0 reference genome were probably due to differences in SNP calling. Differential accessibility in the absence of considerable sequence variation suggests a significant *trans*-contribution; nevertheless, it is well appreciated, and indeed supported by the data presented here, that overall sequence variation of a regulatory site is not a great predictor of function. A single mutational event can suffice to disrupt an essential nucleotide of a transcription factor motif and hence cause altered accessibility (Vierstra et al. 2015; Liu et al. 2014; Stern and Frankel 2013; Kimura et al. 2008). Conversely, as we show, other regulatory sites maintain accessibility despite being hypervariable, implying that sequence variation does not necessarily affect transcription factor binding.

## DISCUSSION

The current challenge in annotating regulatory regions is connecting specific base pairs with regulatory function. We and others have previously attempted to predict the function of regulatory regions by using TF motif content and footprint presence (Segal et al. 2008; Sullivan et al. 2014; Vierstra and Stamatoyannopoulos 2016). There are several limitations to this approach, including TF binding site sequence specificity that differs depending on context (Slattery et al. 2011; Jolma et al. 2015), occupancy that is imperfectly gauged by most current footprinting algorithms, particularly those that do not control for nuclease bias and the particular binding tendencies of different TFs (Vierstra and Stamatoyannopoulos 2016), and TF bindings motifs, especially for large TF families, that can be highly similar. Here, we examined the effect of sequence variation within positionally and presumably functionally homologous regulatory sequence (DHSs) of closely related *A. thaliana* strains to gain insight into regulatory sequence evolution.

As expected, we found that DHSs with changes in both sequence and accessibility across strains are much more likely to reside near genes with different expression than DHSs without these changes. It is well-established that TF-mediated gene regulation is sequence-driven. In addition to the multitude of genetic studies of regulatory regions (Carroll 2008; Wittkopp and Kalay 2011), an elegant large-scale demonstration of this principle comes from a mouse model of Down syndrome in which the extra chromosome 21 copy is a human chromosome 21. The location of TF binding chromosome-wide was primarily determined by the *cis*-component (regulatory sequence) rather than the *trans*-component; *i.e.*, mouse TFs bound the human chromosome 21 in much the same pattern as the homologous human TFs would in a human cell (Wilson et al. 2008; Coller and Kruglyak 2008).

Perhaps more surprisingly, we found that gene regulation can be remarkably robust to sequence perturbations in DHSs; only a quarter of DHSs with changes in both sequence and accessibility resided near genes with altered expression. Furthermore, the majority (hundreds) of DHSs with high sequence variation displayed neither a change in accessibility nor in nearby gene expression. An obvious caveat, however, is our assumption that a given DHS represents the regulatory region for the closest nearby gene. While certainly not perfect, there is no obvious analytical alternative for assigning DHSs to target genes (Sullivan et al. 2015); moreover, extensive *Drosophila* studies suggest that regulatory elements most often target a neighboring gene or, if the regulatory element resides in an intron, the host gene (Kvon et al. 2014). This pattern is also frequently observed for regulatory element-target gene pairs in other organisms (Liu et al. 2014; Stern and Frankel 2013; Vierstra et al. 2015); even when a regulatory element resides very far away from its target gene, in most cases, there are no other genes residing in the intervening sequence (Clark et al. 2006; Stam et al. 2002; Guenther et al. 2014; Kvon et al. 2014).

Assuming correct target gene assignment for the majority of DHSs, the question remains how this regulatory robustness is generated in light of considerable underlying sequence variation. Careful dissections of individual regulatory regions and genome-scale studies implicate several mechanisms for regulatory robustness, including shadow enhancers, transcription factor redundancy and motif redundancy, among others (Payne and Wagner 2015). Nevertheless, the amount of largely unannotated sequence variation in supposedly functional genomic regions among these closely related *A. thaliana* strains was surprising and calls for future sequencing efforts with longer reads. In particular, we observed hypervariable DHSs that were preserved in length among strains, but highly disparate in sequence, showing either a Col-DHS allele or a non-Col-DHS allele. This observation is consistent with a single mutation event that generated the dramatic nucleotide differences between the allelic forms. DHSs that coincided with hypervariable sequence and changed in accessibility predicted expression changes in nearby genes almost as well as DHSs deletions.

Our assessment of the association between gene expression and DHS changes in sequence and accessibility is limited by the finite number of conditions in which expression data were collected (*i.e.* association between gene expression and DHS changes may be condition-specific). Accessibility frequently reflects binding of TFs poised for activation rather than actively driving gene expression in a particular condition (Keene et al. 1981; Sullivan et al. 2014). Deletion of a poised accessible site should not affect expression of a neighboring gene; however, expression of the neighboring gene would be affected in conditions in which the TF is driving its expression. Indeed, we discovered conditional effects of deleted DHSs for several genes, supporting the concept of poised TFs. Sequential cooperative binding of TFs, TF oligomerization, activating cofactors or modifications are necessary to activate poised TFs and drive conditional gene expression (Todeschini, Georges, and Veitia 2014; Slattery et al. 2014).

Finally, and perhaps most surprising, we found that the vast majority of DHSs that differ in accessibility across strains show no underlying sequence variation, at least in the available short-read-derived draft genomes. At a first glance, this result suggests a considerable role of the *trans*-component (*trans*-acting factors that directly or indirectly regulate gene expression) in diversifying gene regulation and phenotype, seemingly challenging the paradigm that evolution largely acts on regulatory regions rather than coding regions. The prevalence of regulatory evolution over coding evolution is thought to arise from the reduced pleiotropy of regulatory changes, the phenotypic effects of which are often tissue-or condition-specific and hence less deleterious, as well as the largely additive effects of regulatory changes that increase the efficiency of selection (Wittkopp and Kalay 2011; Meiklejohn et al. 2014). At this point, the empirical evidence supporting this prediction is overwhelming (Carroll 2008; Wittkopp and Kalay 2011).

We speculate that our finding reflects network effects, as previously suggested (MacNeil et al. 2015; Fuxman Bass et al. 2016). The observed changes in chromatin accessibility without underlying sequence variation are highly unlikely to arise through TF coding changes that alter their DNA binding. Rather, our finding likely reflects the propagation of upstream perturbations in the regulatory network; for example, activity changes in a few upstream ‘master regulator’ TFs may result in lowered abundance or binding of many downstream TFs. This interpretation resonates with the empirical findings of a growing number of studies (Beadle 1972; Doebley and Stec 1991; Studer et al. 2011; Kliebenstein et al. 2006; DeCook et al. 2006; J. Wang et al. 2010; Holloway et al. 2011; Fu et al. 2009; Lemmon et al. 2014).

In summary, the anatomy of regulatory regions appears to be fundamentally more challenging to elucidate than the anatomy of coding regions, from the simple task of mapping them to pinpointing single base pairs of functional consequence. Our understanding of coding sequence anatomy has advanced to the point where we can computationally predict important domains within proteins; indeed, large-scale cross-species analysis of coevolution allows accurate predictions of protein structures, a prerequisite for fully understanding gene function (Ovchinnikov et al. 2015; Tang et al. 2015; Stein, Marks, and Sander 2015). In contrast, although gene regulation is sequence-driven, sequence conservation in regulatory regions is a much less reliable indicator of functional importance than it is for coding sequence.

However, chromatin accessibility is a reliable metric for potential regulatory function of a locus and offers a means to generate useful shortlists of putative regulatory elements (ENCODE Project Consortium 2012). Sites with different levels of accessibility among strains and species are likely enriched for variation that is important for phenotype. Requiring perfect correlation between chromatin accessibility and expression of nearby genes to conclude that a locus is a functional regulatory element would be analogous to requiring a mutant phenotype before concluding a gene is functional — only a tenth of *A. thaliana* genes would pass this stringent definition of functionality after decades of classic genetic analysis (Lloyd and Meinke 2012). Determining the anatomy of accessible regulatory regions remains an essential, albeit challenging, task of genome annotation. Several approaches such as deep mutational scanning of specific regulatory elements (Patwardhan et al. 2009; Fowler et al. 2010) or genome-scale enhancer and promoter screens across species (Arnold et al. 2013) hold promise for solving this challenge, as will new technologies yet to emerge.

## METHODS

### Plant material

Transgenic INTACT lines harboring the UBQ10:NTF construct (Sullivan et al. 2014) and ACT2:BirA construct (Deal and Henikoff 2010) were created for each genetic background using double *Agrobacterium tumefaciens* transformation. Transformants were selected on plates containing BASTA(15 uM) and KAN (50 ug/ml for all strains except Ler-1, which received 15 ug/ml). Transgenic lines are available from ABRC under the following accession numbers: CS68650 (Bay-0), CS68651 (Bur-0), CS68652 (Cvi-0), CS68653 (Est-1), CS68654 (Ler-1), CS68655 (Shah), CS68656 (Tsu-1), CS68649 (Col-0).

### Sample preparation for DNase I-seq and ATAC-seq

Seeds (0.1g) were surface sterilized by treating with 70% EtOH with 0.5% Triton for 10 minutes followed by 5 minutes in 95% EtOH. Seeds were dried completely on sterile filter paper and plated on 150mm petri plates containing 50ml 1XMS with 0.8% agar covered by a sterile #1 filter circle cut to size (Whatman, GE Healthcare UK ltd). Plates were sealed with micropore tape, double wrapped with aluminum foil and stratified for 3 days at 4C. Stratified plates were unwrapped and moved to LD conditions (16hr light, 22°C; 8hr dark, 20°C) in a growth chamber (Conviron CMP5090, Controlled environment ltd. Winnipeg, Manitoba, Canada) and grown for 7 days. Samples were harvested at the same time of day and nuclei were collected and DNase I treated as in Sullivan et al. 2014, Bay-0, Bur-0, Col-0, Est-1 and Tsu-1 DNase I-seq samples were labeled as DS22973, DS23077, DS21094, DS22974, DS22968, respectively. ATAC-seq was performed as in Buenrostro et al. 2013, except transposition of INTACT-purified bead-bound nuclei was performed at 37C for 30 minutes.

### Preparation of ecotype data

Short read whole-genome shotgun data (single and paired-end reads) for Bay-0, Bur-0, Est-1, Tsu-1 and Col-0 were downloaded from the 1001 Genomes project for *A. thaliana* (Gan et al. 2011) http://1001genomes.org/index.html). Single-end DNase I-seq reads were aligned to the appropriate reference genomes using bwa version 0.5.6 with default parameters. ChrC and ChrM reads, and centromeric regions from (Clark et al. 2007) (Chr1:13,698,788-15,897,560; Chr2:2,450,003-5,500,000; Chr3:11,298,763-14,289,014; Chr4:1,800,002-5,150,000; Chr5:10,999,996-13,332,770) were filtered out, and the remaining reads from each sample (Bay-0 n=31,756,881; Bur-0: n=27,556,544; Col-0 n=43,969,877; Est-1 n=30,787,644; Tsu-1 n=17,892,297) were subsampled to 17.5 million reads each. Per-base DNase I cleavages, hotspots (John et al. 2011), and DHSs (peaks) were called on the subsampled data sets as before (Sullivan et al. 2014).

### Identification of differential DHSs

Bay-0, Bur-0, Est-1, Tsu-1 and Col-0 DHSs were merged to create a “union” set of 49,088 DHSs. Per-base DNase I cleavages within each merged DHS were calculated for each ecotype. DNase I cleavages within each DHS were then summed across all five strains. Variable DHSs were then identified based on their coefficient of variation (CV), which is the standard deviation of DHS accessibility across the five strains divided by the mean in DHS accessibility across the five strains. CV was chosen as a metric because the standard deviation of DNase I cut count is a reasonably linear function of the mean DNase I cut count for 83% of DHSs (those with mean DNase I cut count above 63) (see Supplemental Fig. S2). The CV threshold (CV = 0.56) was chosen because all DHSs which are predicted to be affected by a deletion (i.e. DHSs with zero per base DNase I cleavages in at least one ecotype) had a CV greater than or equal to the threshold. This CV threshold corresponds to the top 15 percentile of DHS variability.

To identify DHSs that are differential between Bur-0 and Col-0, we first found the set of (42,656) union DHSs between Bur-0 and Col-0, then identified 4,055 dDHSs from our above five-strain analysis that overlap the Bur-0/Col-0 union set. We made a histogram of the relative mean differences of DNase I cut count within each of those 4,055 dDHSs between Bur-0 and Col-0. Although the relative difference values ranged from −2 to 1.8, a large fraction (3,283; 80%) of these dDHSs had a relative difference of less than −0.3. We therefore defined dDHSs overlapping union Bur-/Col-0 DHSs with a relative difference of less than −0.3 as differential (and higher in Col-0) between Bur-0 and Col-0. We used a similar approach to identify DHSs that are differential between Bay-0 and Col-0 (44,148 Bay-0/Col-0 union DHSs; 4,046 dDHSs overlapping; range of relative differences: [−2, 1.9]; 3363 (83%) differential DHSs between Bay-0 and Col-0).

Browser tracks reflect sliding window histograms of DNase I cut counts with bin size 150 bp and slide size 20 bp. Tracks consist of adjacent non-overlapping bars, where each bar is 20 bp in width and has a height equal to the total number of DNase I cut counts in the larger 150 bp window in which the 20 bp bar is centered.

### Bay-0 draft genome assembly (from D. Weigel)

Bay-0 genomic DNA was isolated from leaf tissue. A paired-end library (PE) was prepared from gDNA sheared to ~500 bp using an S2 Focused-Ultrasonicator (Covaris Inc., MA, USA), and the TruSeq DNA PCR-Free Library Preparation Kit (Illumina, Inc., San Diego, US-CA). A mate pair library (MP) was prepared using the Nextera Mate Pair Library Prep Kit (Illumina, Inc., San Diego, US-CA), following the gel-plus protocol with ~8 kb inserts. The PE library was sequenced on an Illumina MiSeq while the MP library was sequenced on an Illumina HiSeq2000 instrument. The PE data set comprised 28 million read pairs while the MP dataset comprised 43 million read pairs after sequencing. Both datasets were adapter and quality trimmed using skewer (version 0.1.124; parameters -Q30 -q30 -l60). PE data was assembled with DISCOVAR de novo (release 52488, default parameters). Contigs were scaffolded with BESST (version 2.2.0; default parameters) using PE and MP data as input. Scaffolds were validation and fixed using REAPR (version 1.0.17; default parameters) again using PE and MP data as input. The final assembly consisted of 59,594 sequences, totaling 138Mb with a N50 of 1,6Mb.

### Analysis of reference bias

Both Bay-0 and Col-0 DNase I reads were aligned to Bay-0 and Col-0 reference genomes using different alignment stringencies, (i) perfect (maximum edit distance between read and reference of 0; maximum number of gap opens of 0), (ii) default settings (maximum edit distance between read and reference of 0.04 of read length; gap open penalty of 11), (iii) relaxed settings (maximum edit distance between read and reference of 0.06 of read length; gap open penalty of 15). 17.5M aligning reads were retained for each of these 12 alignments. Hotspots were called for each of the 12 alignments. Six sets of union DHSs were called for pairs of alignments with similar alignment settings (eg: perfect) and to the same genome (eg: Bay-0). For example, to obtain one set of union DHSs, we merged the DHSs derived from the default-setting alignment of Bay-0 DNase I reads to Bay-0 and the the DHSs derived from the default-setting alignment of Col-0 DNase I reads to Bay-0. For each of the six sets of union DHSs, we then identified the 1,000 union DHSs in which the number of DNase I cut counts in Bay-0 and Col-0 were most different, using CV as our metric of distances, as above. Within each of these 1,000 most-different union DHSs, we identified whether the union DHS was most open in Bay-0 (orange) or Col-0 (green).

### Variable DHSs near gene families & GO term enrichments

Gene family information was downloaded from TAIR (gene_families_sep_29_09_update.txt). Family members that did not have a corresponding ATG number were filtered out. Overlaps between genes nearest to variable DHSs and family members were tabulated. Hypergeometric tests (phyper in R; q = number of genes nearest to a variable DHS in each family, m = number of genes nearest to variable DHSs in the background, n = number of gene that are not nearest to a variable DHS in the background, k = number of family members in the family) were used to determine the significance of the overlaps between genes nearest to variable DHSs or genes in deletions and family members. GO term, and INTERPRO enrichments were performed using DAVID (Huang, Sherman, and Lempicki 2009). Only the enrichments with FDR less than 0.05 are presented.

### Identification of WGS-called deletions >300 bp in length in the ecotypes

Deletions were identified by calculating the mean x-coverage in all 150 bp sliding windows (overlap = 20 bp) for each ecotype reference using the 1001 genomes (http://1001genomes.org/) short read data. The x-coverage in each window was normalized by dividing by the x-coverage observed for Col-0 short reads mapped back to the reference, which controls for regions of the genome that are more readily sequenced with short reads. We identified putative deletions by taking the 150 bp windows in the bottom 1% of normalized coverage and merged overlapping windows. We then merged putative deletions from different ecotypes that were within 1 kb of each other, to generate the predicted deletions set used in this analysis. Using this method, we determine that 1,975,980 bps, 1,833,441 bps, 2,002,154 bps, 1,712,785 bps are deleted in Bay-0, Bur-0, Est, and Tsu-1, respectively, relative to the Col-0 reference.

### Identification of WGA-called deletions

We aligned the Bay-0 draft genome to the Col-0 reference genome using MUMmer (Kurtz et al. 2004), and defined regions of the Col-0 reference genome that did not have an alignment to the Bay-0 draft genome of 1 kb or more as WGA-called deletions.

### PHRAP-based *de novo* assembly

For each of the 4,508 differential DHS most accessible in Col-0 (dDHS-C), we identified a padded window, encompassing the dDHS-C and including 200 bp extra on either side. We then extracted all reads mapping to that padded window, as well as all mates of reads mapping to these padded dDHS-C windows, regardless of whether the mate mapped to the region. Next, we extracted Col-0 sequence for these padded dDHS-C windows and generated a backbone fasta file with accompanying quality file, setting the quality of each base to zero. After converting all the fastq files to pairs of fasta and fasta.qual files, we used PHRAP to assemble the read pairs and the backbone with relaxed settings (gap extension penalty of 0; minscore of 20; minmatch of 6). The longest contig produced is the patched Bur-0 sequence. Ten of these patched Bur-0 sequences are compared to Sanger sequences and to published sequence (Gan et al. 2011) in Supplemental Table S7. Indicated above each Bur-0/Col-0 alignment is the number of reads used in the PHRAP assembly and the number of reads used to generate the longest contig.

### Proximity of types of DHSs to differentially expressed genes

For each type of DHS, 100 subsamples of 50 DHSs each were drawn. Each sample of 50 DHSs created a distribution of distances to the TSS of the nearest gene with differential expression between Col-0 and Bur-0. For each 50-DHS subsample, the fraction of DHSs within 5 kb up- and down-stream of the nearest gene with differential expression was recorded. The average of these fractions over the 100 50-DHS subsamples is displayed in (Fig. 4A, top). Similarly, for each 50-DHS subsample, the DHS-to-TSS distances in various quantiles [0, 0.02, 0.04, …, 0.98, 1] were recorded. For each quantile we determined the average distance in that quantile among the 100 50-DHS subsamples to get the average quantile values displayed in (Fig. 4B, bottom).

### Conditionally expressed genes are near more DHSs

We used cuffdiff (Trapnell et al. 2013) to identify 1,477, 1,021, 1,312, and 1,365 genes that were differentially expressed (p-value <=0.05) in Col-0 in four different control vs. treatment experiments, respectively: (i) dark-grown seven day-old seedlings compared to dark-grown seven day-old seedlings + 1 day of light exposure (LD, 16h light/8h dark); (ii) LD-grown seven day-old seedlings before and after heat-shock (45C for 30 minutes); (iii) seven day-old seedlings with and without auxin treatment (auxin-treated seedlings were sprayed with 750ul 1uM IAA made from 100mM stock in 70% ethanol, diluted in dH20; no auxin-seedlings were mock sprayed with 750ul 0.0007% ethanol); (iv) dark-grown seven day-old seedlings with and without 5uM brassinazole (BRZ); 7 days in the dark on plates containing 5uM brassinazole (BRZ). All RNA samples are publicly available (http://www.plantregulome.org/public/rna/).

### Response to prolonged UV irradiation

Col-0, Bur-0, *rev3* (CS65883) and *rev7* (SALK_014571C) seeds were germinated and grown for six days on MS Basal Salt plates, under white light (16h light/8h dark). Seven-day old seedlings were then transferred to soil and grown an additional five or seven days under UV irradiation (Zilla, 17W full-spectrum T8 fluorescent bulb). Pools of 32-74 seedlings per genotype were carefully removed from soil and weighed after a total of 12- and 14-days of growth. Fresh weight with and without UV was compared.

### Testing for conditional expression

Bur-0 and Col-0 seedlings were grown on MS Basal Salt plates for six days at 22C, in either constant darkness or long-days (LD, 16h light/8hrs dark). Both dark- and light-grown seedlings were then grown an additional 24hrs in (1) LD 22C, (2) constant darkness 22C, (3) constant white light 4C and (4) constant darkness 4C. RNA was extracted with Trizol (Invitrogen) and oligo-d(T) cDNA generated from 0.2-0.5ug total RNA with the Revert Aid First Strand cDNA synthesis kit (Thermo Scientific). Relative expression levels were determined using qPCR (see Supplemental Table S10 for primers), with the average between Col-0 LD 22C replicates used as baseline. *AP2M* (AT5G46630) was chosen as the reference gene for cDNA input control among several candidates (H. Wang et al. 2014) based on its overall low expression level and lowest coefficient of variation across ecotypes and conditions used (data not shown).

Predicted deletions near these genes were tested using the primers listed in Supplemental Table S10. See Fig. 5C for gel results.

## DATA ACCESS

DNase I-seq data is avaliable at http://www.plantregulome.org/data/releases and at GEO (https://www.ncbi.nlm.nih.gov/geo/; Bay-0: GSM1289371; Bur-0: GSM1289370; Col-0: GSM1289358 and GSM1289359; Est-1: GSM1289372; Tsu-1: GSM1289373). ATAC-seq data have been submitted to SRA.

## ACKNOWLEDGMENTS

This work was supported by grants from the National Science Foundation (MCB1243627 to C.Q. and J.L.N. and MCB1516701 to C.Q.). K.J-B. was supported by a Big Data Training Grant for Genomics and Neuroscience (1T32CA206089-01A1). M.W.D. was supported by an NSF Graduate Research Fellowship. S.F. is an investigator of the Howard Hughes Medical Institute, which supported J.T.C. We thank the Stamatoyannopoulos lab for generously providing sequencing and analysis resources.

## AUTHOR CONTRIBUTIONS

K.L.B., J.L.N., and C.Q. conceived the project.

K.L.B., C.M.A., and C.Q. wrote the manuscript.

A.M.S., J.L.N., D.W. and S.F. provided comments.

K.J-B. and K.L.B. conducted data analysis.

C.M.A., J.R.U., M.W.D., J.C.C., A.M.S., A.A.A., A.T. conducted experiments.

F.B., D.J., and D.W. provided the Bay-0 draft genome.

All authors read and approved the manuscript.

